## Supplementary figures and images for "ESAT-6 undergoes self-association at phagosomal pH and an ESAT-6 specific nanobody restricts M. tuberculosis growth in macrophages"

### Figure 1 - source data 1.png

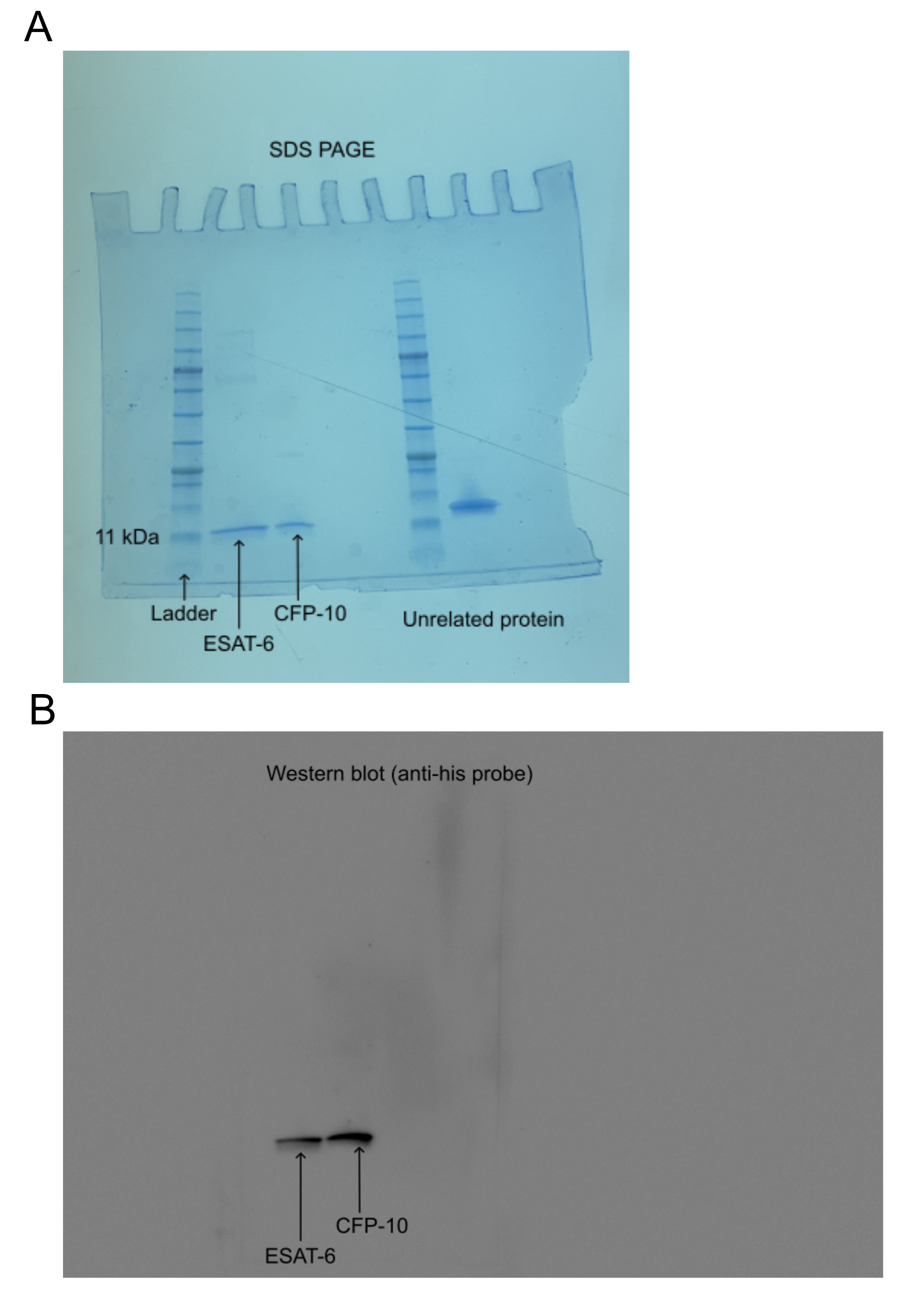

### Figure 1 - source data 1A.png

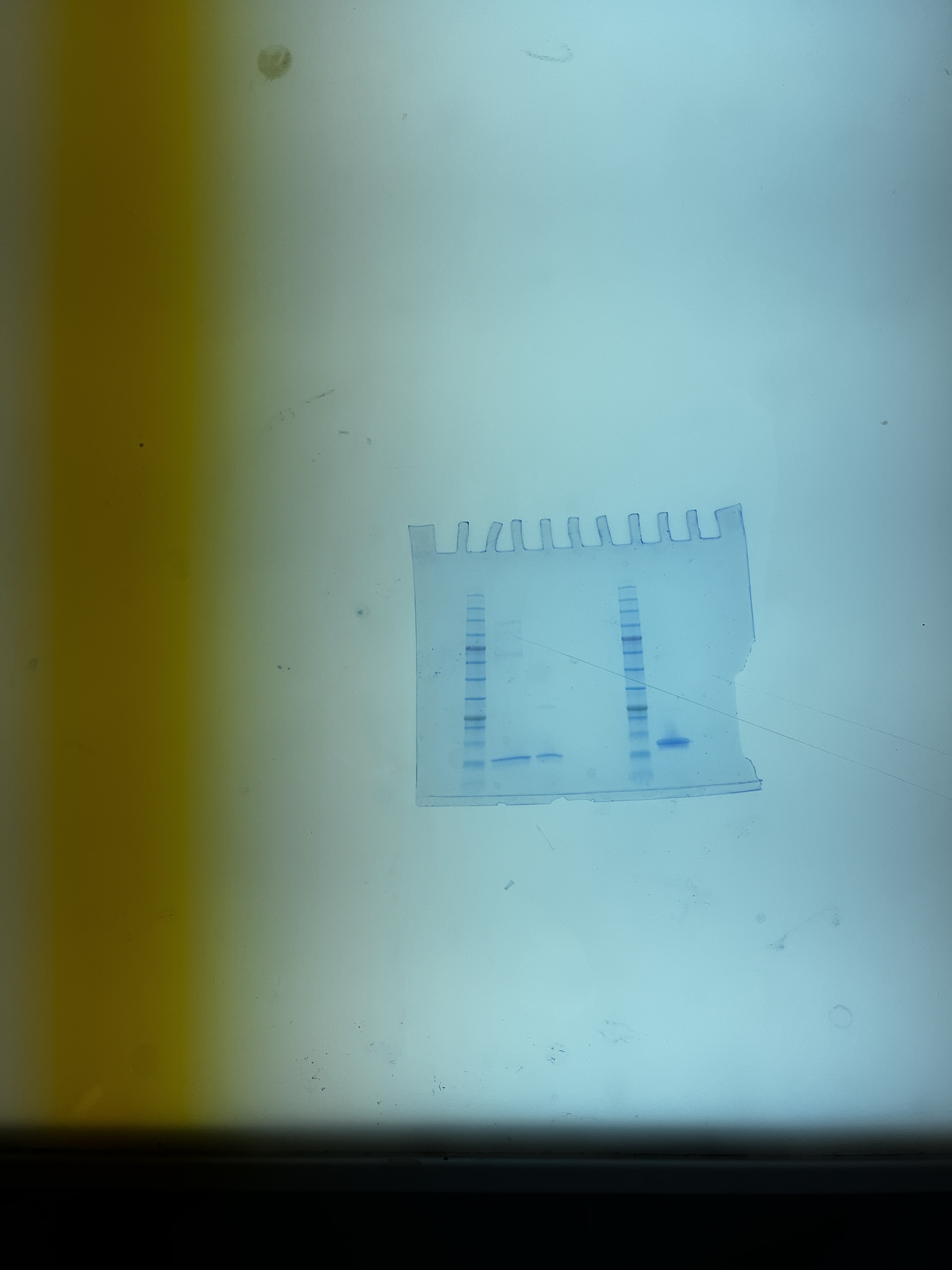

### Figure 1 - source data 1B.tif

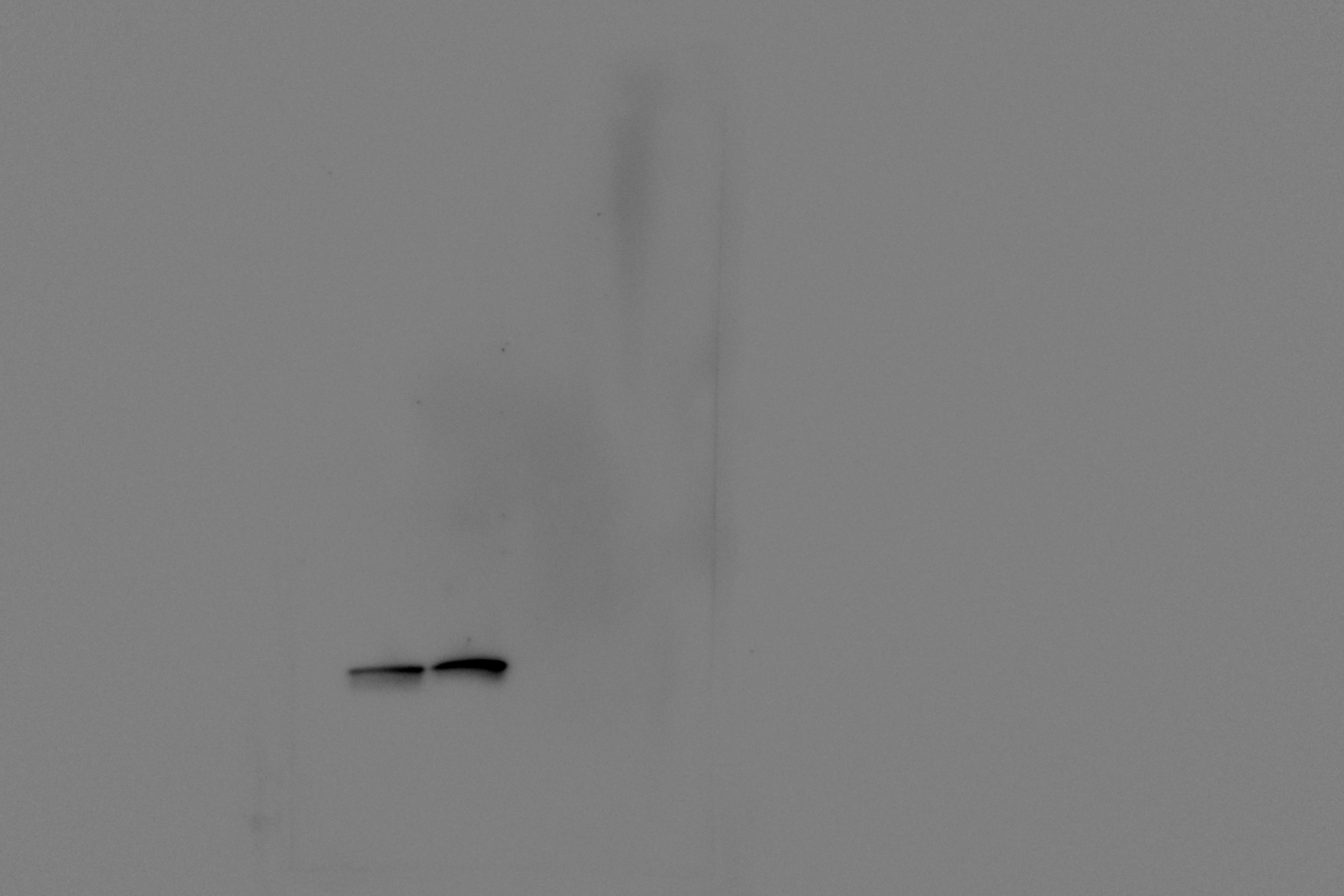

### Figure 2 - source data 1.png

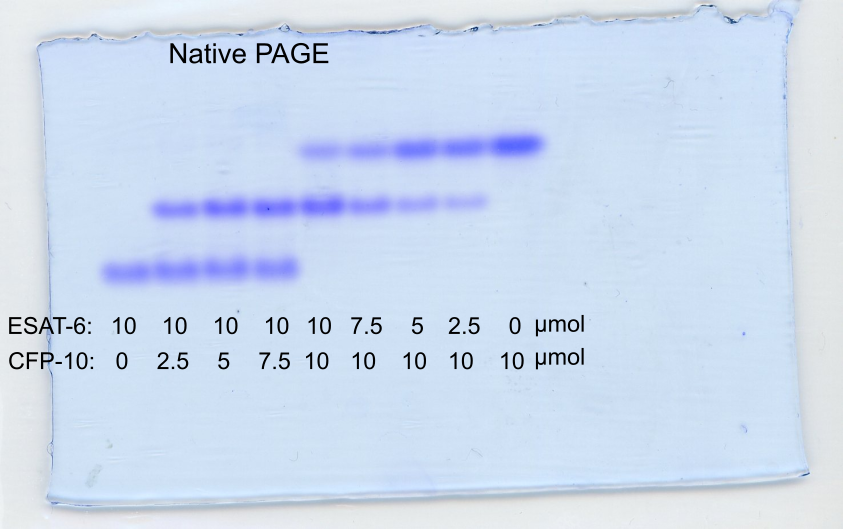

### Figure 2 - source data 1A.jpg

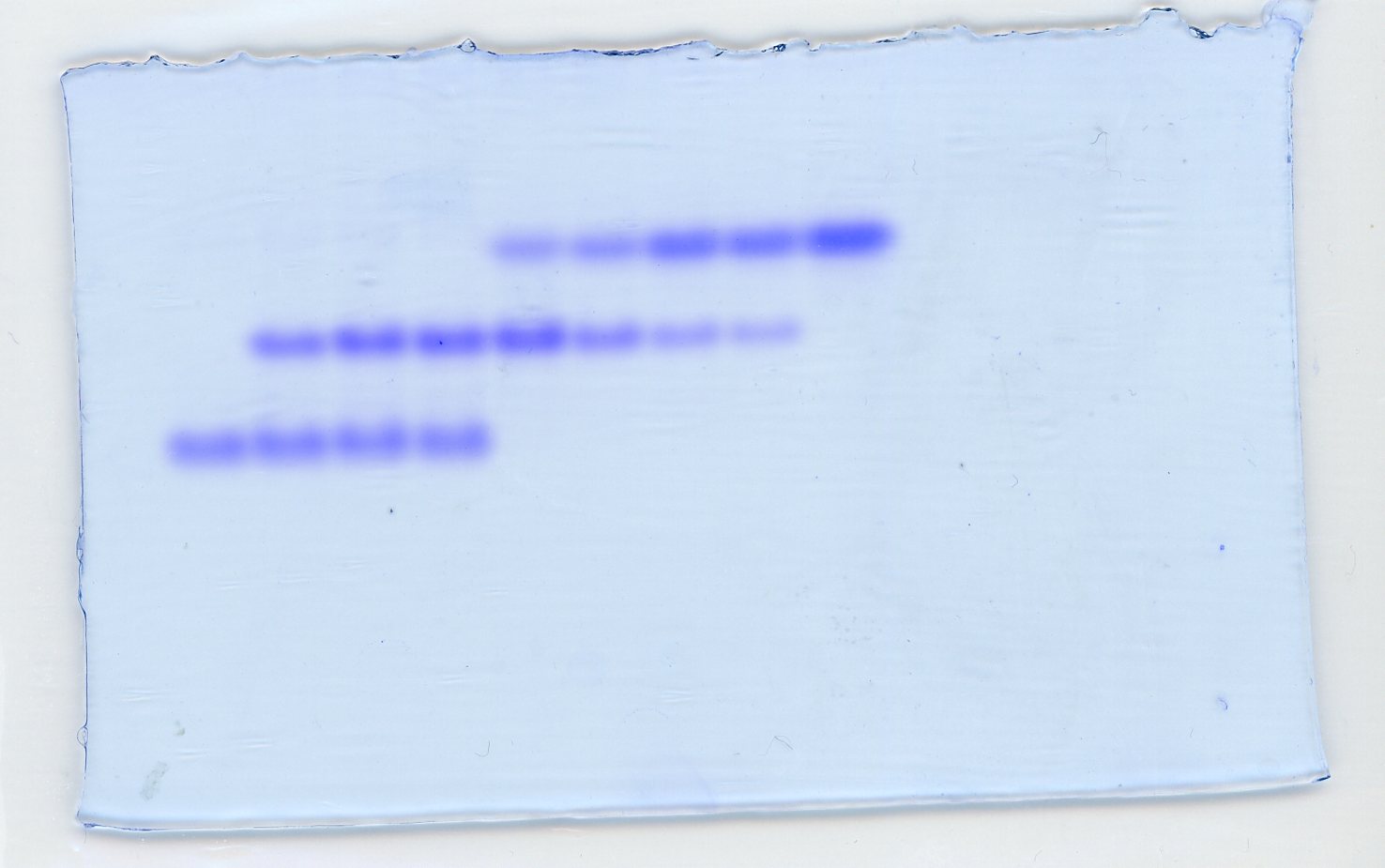
